# Brain criticality emerges with developmental shifts in frequency-specific excitation-inhibition balance

**DOI:** 10.64898/2026.03.26.714563

**Authors:** Andrew Westbrook, Arthur-Ervin Avramiea, Finnegan J. Calabro, Klaus Linkenkaer-Hansen, Shane McKeon, Beatriz Luna

## Abstract

Adolescent brain maturation involves structural changes effecting a shift in excitation/inhibition (E/I) balance, yet the functional implications of these changes remain unclear. One implication is a shift with respect to criticality. Adult brains, at rest, operate near a critical phase transition – at the boundary between an active, excitation-dominant phase, and an absorbing, inhibition-dominant phase. Special properties emerge when neural systems are balanced at criticality, including maximal susceptibility to perturbation, dynamic range, and information transmission. Thus, a clear picture of how adolescent brain maturation affects E/I balance and criticality is needed to understand how maturational processes shape cognition. Here, we leverage the dynamical properties of longitudinally-collected resting-state EEG recordings during N = 310 sessions from 169 healthy human participants ranging in age from 10 to 33 years old to quantify E/I and proximity to criticality. We find that adult brains operate closer to criticality, including spectrally-widespread increases in long-range temporal correlations and amplitude bistability. We also find band-specific changes in excitation versus inhibition whereby the mechanisms driving low-frequency (θ to α) oscillations shift towards lower E/I–possibly because of increasing inhibition–while the mechanisms driving high-frequency (γ) oscillations shift towards higher E/I, possibly because of decreasing inhibition. Opening eyes shifts brains towards lower E/I, and these state-dependent shifts are larger in adults. We simulate developmental effects with a neural mass model of coupled excitatory and inhibitory neurons providing a parsimonious account of how changes in brain dynamics could arise as a function of changes in local connectivity of excitatory and inhibitory neurons. Results indicate developmental movement towards criticality, and greater adaptability to state-specific demands, through adolescence to adulthood, reflecting changes in E/I balance with implications for cognitive development.

## Introduction

At rest, the healthy adult brain operates at a boundary between dynamical regimes [1]. On one side of this phase transition, neural systems are inhibition-dominant and, at the limit, non-responsive. On the other, neural systems are excitation-dominant and, at the limit, chaotic. When excitation and inhibition are balanced near a critical point, emergent dynamics are optimal for computation. They confer susceptibility [1–4] – the propensity for the system to respond to inputs – and scale-free activity – implying persistence of information [5,6] and maximal dynamic range [7,8]. Thus, excitation-inhibition (E/I) balance determines proximity to criticality and sets the stage for diverse cognitive capacities.

Adolescent brains go through numerous changes in cortical architecture including pruning of excitatory synapses, the maturation of inhibitory interneurons, and the expression of GABA receptors, all suggesting a shift in the balance of excitation to inhibition [9–12]. Evidence of a developmental shift in E/I includes a reduction in glutamate relative to GABA metabolites in dorsolateral prefrontal cortex, as measured by magnetic resonance spectroscopy [13,14]. Broader shifts in E/I are also evident in systematic changes in aperiodic power as measured by MEG and EEG [13,15–17]. Namely, with development, oscillatory power is lower (lower offset of a model fit to 1/f power spectra) and power is also reduced by a greater degree in low versus high frequency bands (shallower 1/f slope) in adults. Machine learning and artificial neural network models fit to spontaneous resting-state dynamics in fMRI data also converge on the conclusion that development is associated with widespread reductions in E/I, across the cortex [18,19]. Distinctions across cortical regions in these models furthermore indicate that the rate of change in E/I differs between higher-order association areas, and lower-order, sensorimotor cortex, implying a diversity in the effects of development on excitation versus inhibition across the brain. Interestingly, a developmental shift in E/I in the association cortices, in particular, is linked to IQ in children [20].

A shift in E/I across the brain implies systematic shifting with respect to criticality. Indeed, as adolescents age into adulthood, their brains appear to operate closer to a critical point. Long-range temporal correlations – reflecting persistence and the capacity for integration – are one emergent property of criticality that has been found to grow stronger with development [21]. Signal complexity is another emergent property that is greater at criticality and has also been found to increase from early childhood into adolescence [22]. While long-range temporal correlations and complexity are maximal in critical systems, they diminish under either sub-critical, inhibition-dominance, or super-critical, excitation-dominance [23]. Thus, while brains appear to operate closer to criticality with age, the regime in which youths’ brains operate is not specified by these results. These patterns also do not reveal the direction (increases or decreases in E/I) with which brains shift during development. Finally, there is likely to be a diversity of developmental effects on brain criticality in specific circuits. Even if there is a systemic shift in E/I, whether specific circuits shift towards or away from criticality depends on whether they start in inhibition-dominant, or excitation-dominant regimes.

In this study, we analyze spontaneous brain dynamics in EEG recordings in a sample of 169 healthy human participants ranging in age from 10 to 33 years old assessed longitudinally at approximately 18 month intervals (for a total of 331 recordings) during eyes-closed and eyes-open rest. We derive indices of brain criticality including long-range temporal correlations and bistability, and indices of E/I derived from the dynamics and distributional properties of signal amplitude. Importantly, we compute these metrics spectrally, i.e. across frequency bands, to detect individual differences and age-related shifts in both E/I values and criticality by band [24]. We find that adolescent development is associated with systemic shifting towards criticality. We also find that this shift corresponds with a diversity of changes in E/I metrics in different frequency bands, suggesting a diversity of developmental effects on E/I in distinct brain systems. In a contrast of eyes-closed versus eyes-open rest, we find that opening one’s eyes consistently shifts brains in the direction of stronger inhibition but that this positions some participants’ brains closer to, and others farther away from criticality. We also find that the effect of eyes opening on critical dynamics is larger in adults than in children. Together these results suggest that adult brains both operate closer to criticality with eyes closed and shift more dynamically, with respect to criticality, as a function of processing demands. Finally, we use simulations of an artificial neural network model – which passes through criticality when excitation and inhibition are balanced – to show that we can recapitulate developmental effects on spontaneous EEG dynamics, in each canonical frequency band, by manipulating the relative connection density for excitatory versus inhibitory neurons.

## Results

### Spectrally-widespread strengthening of critical dynamics with age

To test the hypothesis that brains develop towards criticality from adolescence to adulthood, we tested for two emergent properties of a complex, dynamical system at a critical point: long-range temporal correlations, and bistability. Long-range temporal correlations (LRTC) arise in the brain because, when excitation and inhibition are in balance, the effect of a perturbation neither diverges chaotically due to over-excitation nor dies off due to over-inhibition [23]. Bistability arises because critical systems, by definition, operate at the boundary between functional regimes and, therefore, can exhibit properties of both regimes over time [25]. In EEG data, we quantify LRTC in terms of the persistence of fluctuations in band-limited amplitude, using detrended fluctuation analysis (DFA; [26,27] see Methods). Bistability is quantified in terms of the degree to which the amplitude distribution of an EEG trace is best modeled by two distinct modes (bistability, BiS [25,28]), indicating vacillations between dynamical phases, rather than a single mode, which would indicate a single dynamical phase.

As predicted, we find that both spectrally-widespread LRTC and BiS strengthen with age (Figure 1) indicating a developmental shift towards criticality. Mixed effects regressions including both eyes-open and eyes-closed conditions revealed a main effect of age on increases in DFA and BiS across nearly all frequency bands, except for γ where DFA and BiS decrease with age. For example, in the α band (averaged from 8.3–13.4 Hz) DFA exponents (*β* = .351, *p* = 10^-8^) and BiS scores increase with age (*β* = .251, *p* = 10^-5^). Conversely, in the γ band (27.6–57.6 Hz) both DFA (*β* = -.197, *p* = .00176) and BiS decrease (*β* = -.163, *p* = .00981). Here, to account for the apparently non-linear effects of Age, our models are fit to −1/*Age* [29]. These patterns provide evidence that the mechanisms driving low (θ, α, and β) but not high-frequency oscillations (γ) operate closer to criticality with development.

**Figure 1.**
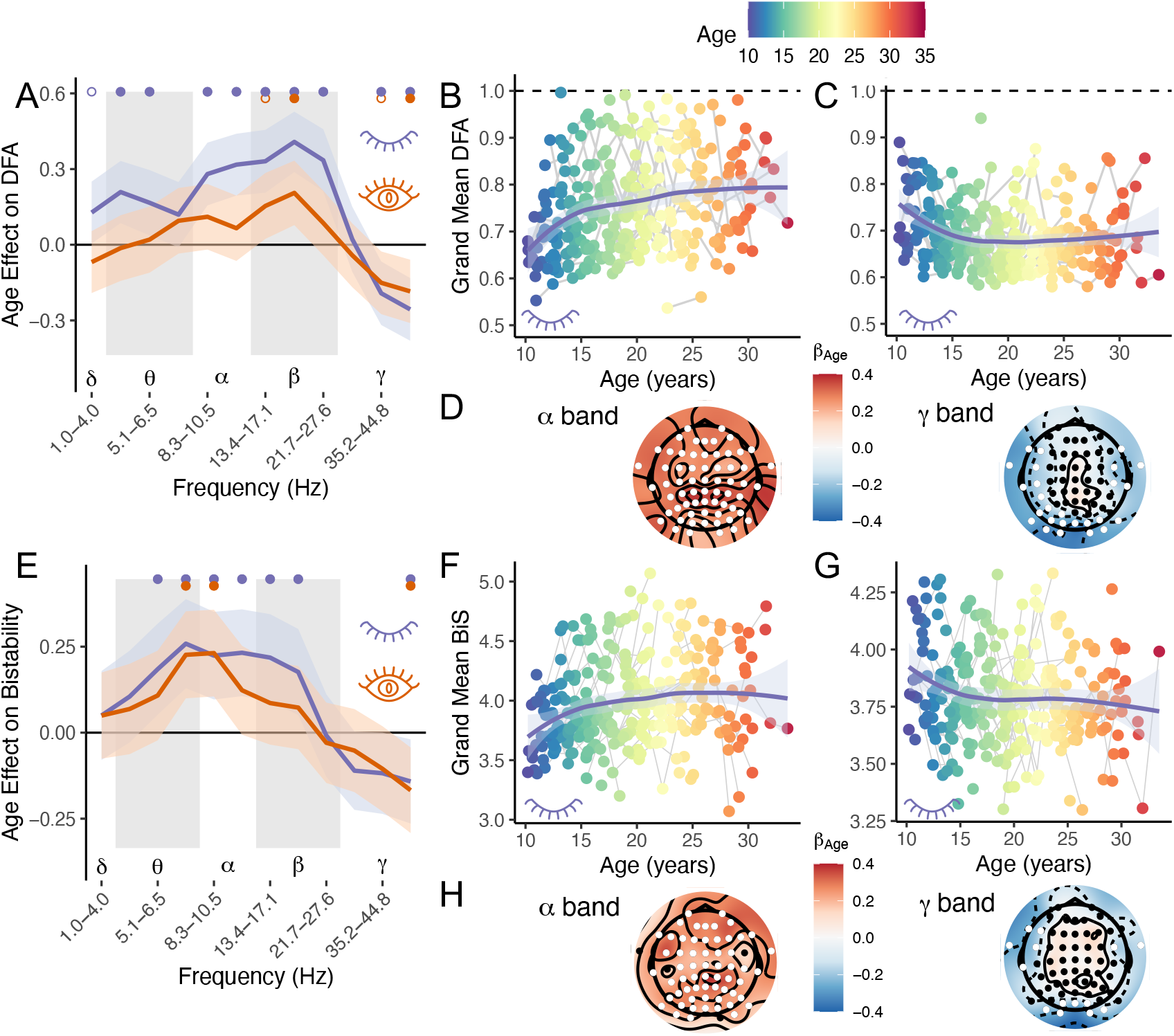
DFA exponents and BiS statistics show spectrally-widespread convergence towards criticality with development. A) Estimates of the fixed effect of age on mean, whole-brain DFA exponents for logarithmically spaced frequency bands from 1.0 – 57.6 Hz in mixed effects models with random intercepts for participants. Orange for eyes-open and purple for eyes-closed. Error bands indicate 95% CI. Closed dots indicate *p_FDR_* < .05, with FDR correction for the number of frequency bands and open circles indicate uncorrected *p_unc_* < .05. B,C) Mean, whole-brain DFA plotted as a function of age for the α band (8.3–13.4 Hz; left panel) and γ band (27.6–57.6 Hz, right panel). A dashed line indicates the theoretical expected value of a DFA exponents when the system operates at criticality (DFA = 1.0). Error bands indicate 95% CI. D) Corresponding, sensor-level correlations between age and DFA exponents in the α and γ bands. White circles indicate sensors where the effect of age was significant at *p* < .01. E) Estimates of the fixed effect of age on mean, whole-brain BiS estimates across frequency bands. F,G) Mean, whole-brain BiS estimates as a function of age in the α and γ bands. H) Corresponding, sensor-level correlations between age and BiS estimates in the α and γ bands.

We also find strong effects of opening eyes on indices of criticality in our mixed effects regression models which included random intercepts for subjects, along with fixed effects for Condition (eyes-open versus -closed) and the Age × Condition interaction as predictors. Both DFA and BiS in the α band dampen when eyes open (DFA: *β* = -.467, *p* = 10^-14^; BiS: *β* = -.179, *p* = .00455) indicating that opening eyes shifts α-generating systems away from the critical point. In contrast, opening eyes strengthens DFA statistics in the γ band (DFA: *β* = .147, *p* = .0180; BiS: *β* = -.0739, *p* = .340). Finally, we note a significant Age × Condition interaction for DFA (in the α band only: *β* = -.302, *p* = 10^-7^; this interaction was not significant in the γ band) suggesting that the effect of development on DFA in the α band is stronger when eyes are closed. At the sensor level, positive effects of Age on both DFA and BiS are widespread, showing up across the surface of the scalp. In contrast, the negative effects of Age on γ-band DFA are more spatially selective, showing up specifically in lateral and posterior electrodes (Figure 1D).

### Age-related changes in the excitation-inhibition ratio

Age-related increases in critical dynamics are consistent with the hypothesis that adolescent development systematically alters the ratio of excitation to inhibition (E/I) – a control parameter which determines proximity to criticality in the brain. The direction of a shift in E/I, however, is not unambiguous from changes in long-range temporal correlations or amplitude bistability, because they decrease when either excitation or inhibition dominate. To determine how development alters E/I, we examine three indices: the functional E/I index (fE/I [30]), which quantifies E/I in terms of correlations between signal amplitude and amplitude fluctuations (positive in subcritical regimes and negative in supercritical regimes), the high-to-low amplitude proportion (E/I_HLP_ [28]), which quantifies E/I in terms of the relative proportion of time in which system with bistable amplitude operates in a high-amplitude, excitation-dominant state versus a low-amplitude inhibition-dominant state, and the branching ratio (σ [31]) which evaluates the rate of growth of neuronal avalanches – or, in EEG data, high-amplitude bursts of neural activity clustered together in time. As E/I increases, avalanches grow faster, reflected in higher branching ratios.

The indices provide converging evidence for frequency-specific developmental effects, with decreasing E/I in mechanisms driving low-frequency oscillations and increasing E/I in the mechanisms driving high-frequency oscillations (Figure 2). Specifically, mixed effects models including Age, Condition and their interaction reveal that θ band (4.0–8.3 Hz) fE/I (*β* = -.223, *p* = 10^-4^) and E/I_HLP_ scores decrease (*β* = -.243, *p* = 10^-4^) with Age. Conversely, in the γ band (27.6– 57.6 Hz) both fE/I (*β* = .261, *p* = 10^-5^) and E/I_HLP_ increase (*β* = .220, *p* = 10^-4^). Interestingly, this pattern suggests that development has a mix of effects on E/I across brain mechanisms associated with distinct frequency bands. There is also evidence of spatial heterogeneity in these patterns (Figure 2D,H).

**Figure 2.**
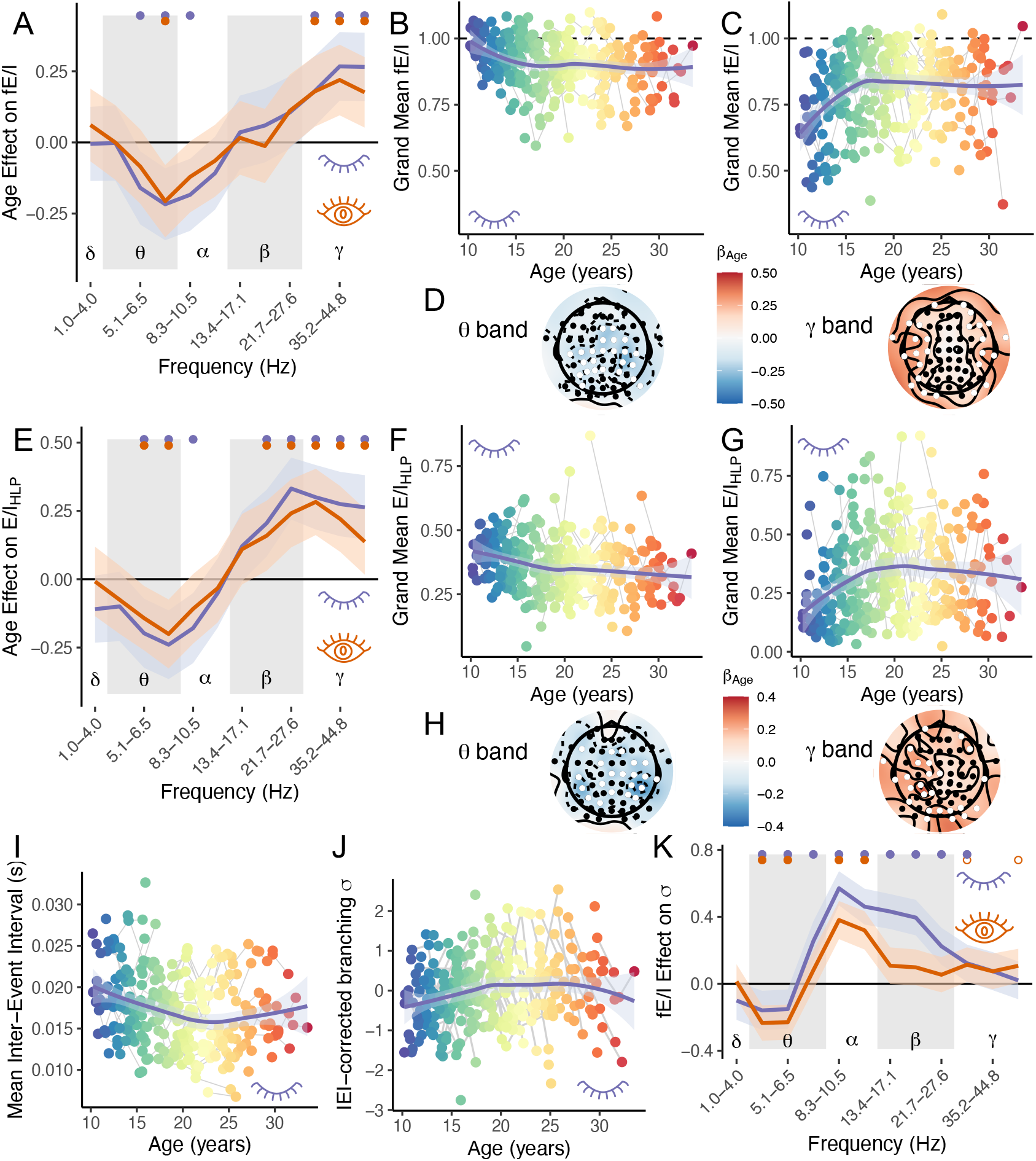
E/I indices fE/I and E/I_HLP_ and the branching ratio all show spectrally-widespread changes with development. A) Estimates of the fixed effect of Age on mean, whole-brain fE/I exponents for logarithmically spaced frequency bands from 1.0–57.6 Hz in mixed effects models with random intercepts for participants. Orange for eyes-open and purple for eyes-closed. Error bands indicate 95% CI. Closed dots indicate *p* < .05, with FDR correction for the number of frequency bands and open circles indicate uncorrected *p* < .05. B,C) Mean, whole-brain fE/I plotted as a function of Age for the θ band (4.0–8.3 Hz; left panel) and γ band (27.6–57.6 Hz, right panel) with eyes closed. A dashed line indicates the theoretical expected value of fE/I exponents when operating at criticality (1.0). Error bands are 95% CI. D) Corresponding, sensor-level effect estimates for Age on fE/I in the θ and γ bands. White circles indicate sensors where the effect was significant at *p* < .01. E) Age effect estimates on mean, whole-brain E/I_HLP_ across frequency bands. F,G) Mean, whole-brain E/I_HLP_ as a function of Age in the θ and γ bands. H) Corresponding, sensor-level effect estimates for Age on E/I_HLP_ in the θ and γ bands. I) Mean inter-event-intervals (IEI) in seconds decrease with Age. J) Controlling for IEIs, the branching ratio increases with Age. K) Estimates of the fixed effect of fE/I in each frequency band on branching ratio values, controlling for Age for both the eyes-closed and -open condition.

The branching ratio (σ) quantifies a global index of the E/I ratio by computing the rate at which high-amplitude EEG bursts (“events”) lead to subsequent bursts, clustered together in time (“avalanches”) [32–35]. When excitation dominates, high amplitude bursts produce subsequent bursts at a higher rate, and the branching ratio is higher. We first observed that mean inter-event intervals decrease with development (Figure 2I). That is, high-amplitude bursts occur more frequently as individuals age. Branching ratio estimates depend strongly on the width of the bin used to discretize timeseries (the mean inter-event interval [32,33]), thus we controlled for mean inter-event-intervals when testing for an effect of Age on branching ratios. A mixed effect regression of branching ratio onto Age (*β* = .154, *p* = .00414) and Condition (*β* = -.217, *p* = 10^-4^), correcting for inter-event-intervals reveals that branching ratios increase with Age (Figure 2J) and decrease, controlling for Age, when eyes open. Thus, with development, high-amplitude events occur more frequently and avalanches grow faster, indicating an increase in global E/I.

To further interpret individual differences in the branching ratio in terms of frequency-specific E/I, we regressed branching ratios onto fE/I, Condition and the fE/I χ Condition interaction, in each of our frequency bands, separately. Because Age has effects on both fE/I and branching ratios, we included Age (as above, −1/*Age*) as a covariate. Our analyses reveal that higher E/I in the α, β, and γ range all predict higher global branching ratios, as expected (Figure 2K). Interestingly, higher fE/I values in the θ range predict lower branching ratios, suggesting that mechanisms which generate θ oscillations suppress high-amplitude EEG bursts.

These indices, considered together with long-range temporal correlations and bistability (Figure 1) reveal that mechanisms that generate low-frequency oscillations move closer to criticality via decreasing E/I, suggesting that low-frequency oscillatory systems move from super-critical to near-critical with development. In contrast, mechanisms which generate high-frequency (γ) oscillations show increasing E/I – helping to account for an increase in branching ratios. Although long-range temporal correlations and bistability are weaker in γ band activity, this does not necessarily imply that high-frequency oscillatory systems move away from criticality. Indeed, fE/I scores (Figure 2A,C) suggest that γ mechanisms begin in sub-critical states early in adolescence and move closer to criticality with development. Thus, developmental changes other than E/I ratios may account for the suppression of long-range temporal correlations and bistability in γ-generating mechanisms with age (see Discussion).

### Age effects on excitation versus inhibition

While the previous analyses quantified the effects of development on the ratio of excitation to inhibition (E/I), we also leveraged orthogonal information provided by an index of combined excitatory and inhibitory strength (E+I) to make inferences more specifically about separable effects of development on excitation versus inhibition. Specifically, we calculated the high- and low-power separation index (E+I_HLS_ [28]), which quantifies combined strength of excitation and inhibition in terms of the difference in power for high- and low-amplitude oscillations. The intuition behind this measure is that when both excitation and inhibition are strong, the swing between high and low amplitude dynamics is larger, as compared to when both are weak.

Mixed effects regressions of E+I_HLS_ onto Age, Condition, and their interaction reveal increases with Age in low-frequency systems (θ, α, and β bands) and decreases with Age in the high-frequency γ range (Figure 3A–D). Thus, combined excitatory and inhibitory drive (E+I) strengthens with development in the mechanisms generating low frequency oscillations, while it attenuates in mechanisms generating γ oscillations. Taken together, information about E/I and E+I values, allow us to infer that inhibitory signaling must increase (or at least increase more than excitation), in low-frequency systems, to account for both a decrease in E/I and an increase in E+I, while it must decrease (more than excitation), in high-frequency systems, to account for both an increase in E/I and a decrease in E+I.

**Figure 3.**
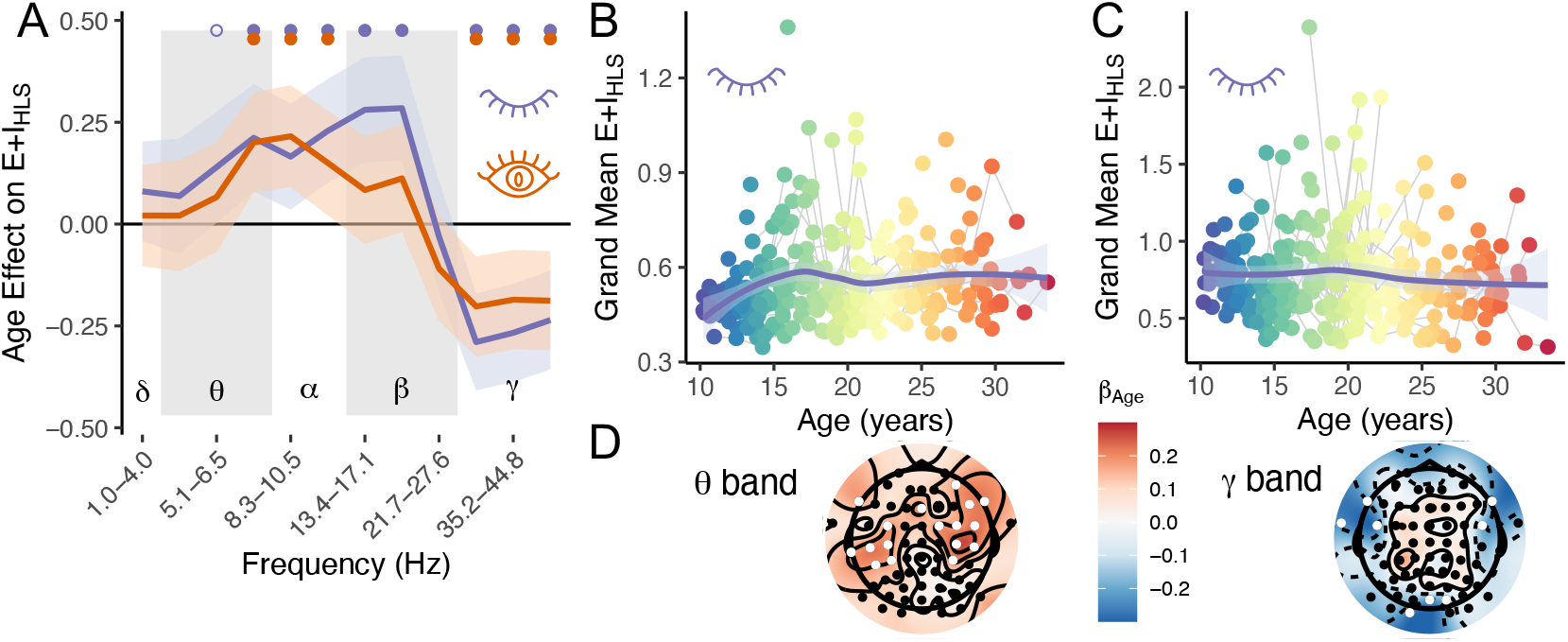
Age changes combined excitatory and inhibitory signaling as measured by the high-to-low power separation index (E+I_HLS_) A) Fixed effect estimates of Age on mean, whole-brain E+I_HLS_ for logarithmically spaced frequency bands from 1.0–57.6 Hz in mixed effects models with random intercepts for participants. Orange for eyes-open and purple for eyes-closed. Error bands indicate 95% CI. Closed dots indicate *p* < .05, with FDR correction for the number of frequency bands and open circles indicate uncorrected *p* < .05 B,C) Mean, whole-brain E+I_HLS_ as a function of Age in the θ and γ bands in the eyes-closed condition. D) Corresponding, sensor-level effect estimates for Age on E+I_HLS_ in the θ and γ bands. White circles indicate sensors where the effect was significant at *p* < .01.

### Opening eyes shifts E/I ratios towards subcriticality

In general, correlations between age and markers of criticality and E/I are slightly, and at some frequencies significantly, stronger with eyes-closed than with eyes-open (Figure 1A,E & 2A,E). The direct effects of opening eyes on E/I and criticality indices, by contrast, are spectrally-widespread. Namely, when eyes are open, there is a systematic shift towards lower E/I (Figure 4C) and a suppression of long-range temporal correlations (Figure 4H), implying a shift to sub-criticality. Regarding E/I, opening eyes decreases branching rates *σ* (t(283) = -10.4, *p* < 10^-16^; Figure 4B versus 4A), and also decreases, e.g., α band fE/I (t(560) = -9.40, *p* < 10^-16^; 4A,B,C) and E/I_HLP_ (t(297) = -8.72, *p* < 10^-16^), within individuals. The decrease in E/I is spectrally widespread, occurring at all frequencies except in the θ band (Figure 4C). The topography of the eyes opening effect on fE/I varies by frequency, with the effect being present across all channels in the α band but restricted to lateral and posterior sensors in the γ band (Figure 4E).

**Figure 4.**
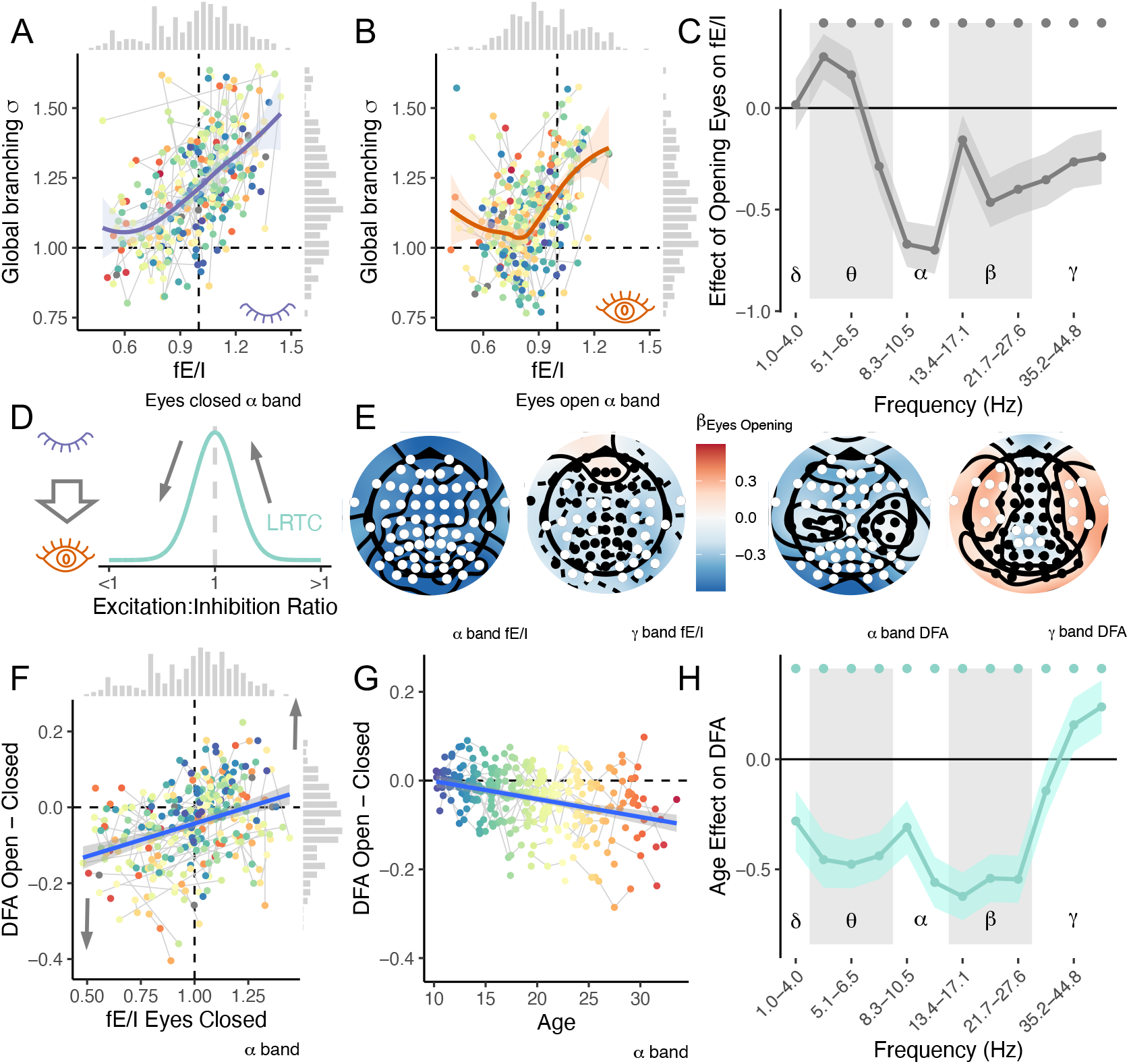
Opening eyes reduces E/I. A) Branching statistic *σ* correlates with fE/I for eyes closed resting data computed in the α band (8.3-13.4 Hz). Dashed lines indicate theoretical values of *σ* and fE/I at criticality. Grey bars show marginal histograms. B) Branching statistic *σ* correlates with fE/I for eyes-open resting data. C) The effect of opening eyes on mean fE/I scores across sensors from a mixed effects regression controlling for Age and the Age × Condition interaction. Solid dots indicate p_FDR_ < .05. D) Conceptual schematic of the effect of opening eyes anticipating a systematic shift towards lower E/I and thereby a shift towards criticality for supercritical brains, and a shift away from criticality for subcritical brains. E) Topography reflecting the effect of opening eyes on, from left to right, α band fE/I, γ band fE/I, and α and γ band DFA scores from a linear regression controlling for participant age and the age × eyes interaction. White sensors indicate *p* < .01. F) The difference in DFA estimates of long-range temporal correlations for eyes-open versus eyes-closed rest plotted against α band fE/I values at eyes-closed. Dark grey arrows show predictions for the difference score to the right and left of the critical value (fE/I = 1.0). G) The difference in DFA estimates for open versus closed eyes plotted as a function of age. H) The effect of opening eyes on mean DFA scores across sensors from a mixed effects regression controlling for Age and the Age × Condition interaction.

Because E/I controls proximity to criticality, the effect of eyes opening on fE/I also implies an effect on brain criticality. Indeed, opening eyes also results in spectrally-widespread decreases in DFA, e.g: in the α band (8.3-13.4 Hz; *t*(298) = -7.37, *p* = 10^-12^). DFA decreases when eyes are open for all frequencies except the γ band, where long-range temporal correlations get stronger (27.6– 57.6 Hz; t(298) = 2.89, *p* = .00412; Figure 3H). The topography of the effect of opening eyes on DFA also varies by frequency, with the effect being widespread across anterior and posterior channels, but not medial channels in the α band. In contrast, increases in γ-band DFA are focal on lateral-frontal sensors (Figure 3E). Interestingly, the effect of opening eyes is stronger with age. Controlling for the effect of fE/I on DFA values, we find a negative effect of Age (α band: β = -.158, *p* = .0138, Figure 3E; β band: *β* = -.328, *p* = 10^-7^; all other bands p ≥ .382) reflecting the fact that adults show a greater state dependence, relative to adolescents and children, with a bigger reduction in long-range temporal correlations when they open their eyes.

Although opening eyes decreases long-range temporal correlations for most frequencies at the group level (Figure 3G,H), the specific effect, at the level of the individual, should depend on E/I balance at baseline. Namely, brains operating in the subcritical regime when eyes are closed should shift away from criticality when eyes are opened, while brains operating the supercritical regime should shift towards criticality when eyes are opened (Figure 3D). Consistent with this prediction, we find that fE/I in the α band at eyes-closed rest correlates positively with the change in DFA exponents when eyes are opened (*β* = .300, *p* = 10^-7^; Figure 3F). This finding confirms a core prediction about long-range temporal correlations as an emergent dynamical property when excitation and inhibition are balanced at criticality and fE/I as a valid measure of individual differences in the E/I control parameter with respect to a critical point.

### Simulating developmental effects by frequency band

Developmental effects on E/I can be recapitulated by an artificial neural network [23,24,28], as a function of excitatory and inhibitory connectivity. The model comprises coupled excitatory and inhibitory neurons (Figure 5B) with varying degrees of local connection density (Figure 5A). Emergent dynamical properties indicate that the model undergoes a critical phase transition, when excitatory and inhibitory connectivity are balanced (black solid line, Figure 5C,D), including strong long range temporal correlations, power-law neuronal avalanches [23], and bistable amplitude distributions [24,28] (Supplemental Figure S1). Importantly, E/I_HLP_ and E+I_HLS_ [28] increase along non-parallel vectors spanning the state space of possible models (Figure 5C,D), making it possible to project developmental trajectories.

**Figure 5.**
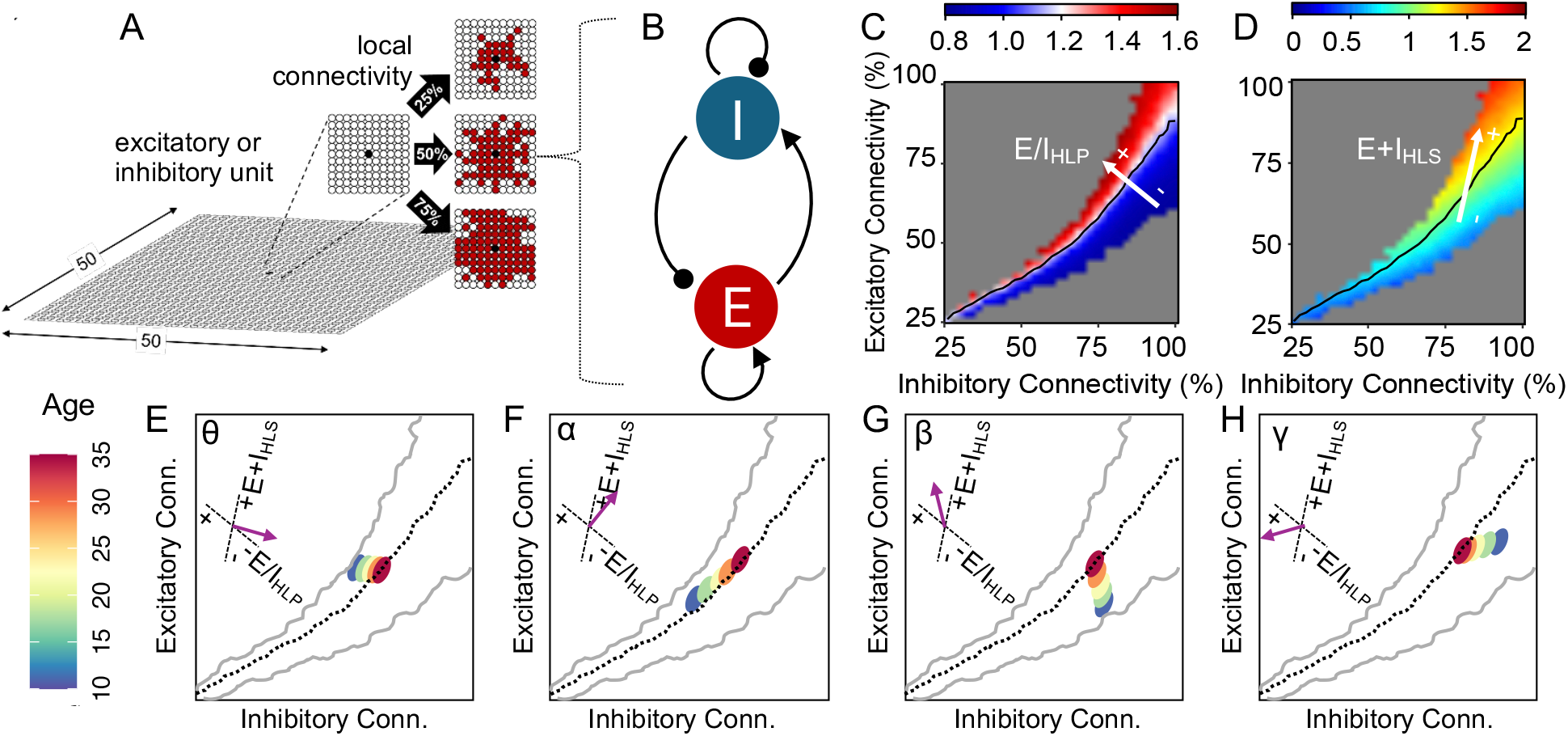
Simulations of an artificial neural network (introduced by [23], using parameters from [28]) recapitulating developmental effects on E/I balance and criticality. A) Network architecture includes coupled, recurrent excitatory and inhibitory units, in a large array (of 50 × 50 units) with local connection density as a model parameter. B) The circuit montage. C) E/I_HLP_ quantified from the amplitude distribution of summed network activity indicates a shift between high amplitude-dominant super-critical dynamics and low amplitude-dominant subcritical dynamics when excitation and inhibition are balanced at criticality, marked by a black line. Red-blue color scale indicates E/I_HLP_. White arrow indicates the direction of the steepest gradient of increasing E/I_HLP_ in the phase space. C) E+I_HLS_ quantified from summed network activity. Rainbow color scale indicates E+I_HLS_. White arrow indicates increasing E+I_HLS_ in the phase space. E-H) Feasible developmental trajectories (from ages 10, in red, to 35 in blue) across the phase space, for mechanisms driving θ, α, β, and γ oscillations based on E/I_HLP_, E+I_HLS_, and fE/I values. Inset shows the empirical direction of each trajectory, indicated by a purple arrow, based on empirical E/I_HLP_ and E+I_HLS_ values.

We illustrate plausible developmental trajectories for mechanisms associated with each of the canonical frequency bands (θ, α, β, and γ), in terms of excitatory and inhibitory connection density (Figure 5E–H). The approximate position of the start (blue, Age = 10) and end (red, Age = 35) of each trajectory, relative to criticality (black dotted line) was determined by taking into account developmental changes in long-range temporal correlations and bistability (for determining proximity to criticality; c.f. Supplemental Figure S1), fE/I (for determining the sub-versus super-critical start and end points), and E/I_HLP_ and E+I_HLS_ (for determining direction vector). The mean direction of each trajectory was determined based on developmental effects on E/I_HLP_ and E+I_HLS_ values which span the state space [28]. While multiple mechanisms could effect an E/I shift, these simulations reveal that changes in excitatory and inhibitory connection density are sufficient to recapitulate developmental changes in spontaneous, resting EEG dynamics observed in our sample.

## Discussion

Age-related changes in excitatory versus inhibitory signaling capacity suggest that the brain systematically shifts towards criticality, with development, from adolescence to adulthood. Consistent with this hypothesis, we replicate prior evidence that long-range temporal correlations grow stronger from adolescence into adulthood (Figure 1) [21]. We also find greater bistability in amplitude distributions. Furthermore, we find developmental effects across the frequency spectrum, but with variations by band. Namely, while markers of criticality become stronger in low-frequency bands (θ-through-β), they become weaker in the γ band.

Our findings also provide new insights into the changes driving shifts in critical dynamics. Namely, we find frequency-specific changes in functional E/I including decreases in excitation versus inhibition in the θ-through-α range and increases in high-β and γ according to complementary measures: fE/I and E/I_HLP_ (Figure 2). This developmental pattern converges with prior evidence of shallower slopes in the aperiodic spectrum with age, reflecting relatively higher power in fast versus slow oscillations [13,15–17]. Interestingly, we also find that branching ratios increase and that high amplitude bursts occur more frequently, suggesting an overall increase in the E/I ratio with development. Further analyses reveal that individual differences in branching ratios, controlling for age, correspond to higher E/I values in high frequency systems (α-through-γ) and lower E/I values in the θ band. Collectively, these results suggest a developmental increase in E/I in high-frequency systems and a developmental decrease in E/I in low-frequency systems, with consequences for the balance of spontaneous activity across the frequency spectrum.

Decreases in E/I indices for θ-through-α support the hypothesis that developmental pruning of excitatory synapses and maturation of inhibitory interneurons decreases the E/I ratio, especially in higher-order association areas [10,11,36]. Convergent evidence comes from analyses of prefrontal MR spectroscopy data showing decreased glutamate versus GABA metabolites with development [13,14]. Similar support comes from analyses of slow (fMRI BOLD) brain dynamics. For example, whole-brain simulations of reciprocally-connected excitatory and inhibitory populations best explain developmental changes in slow BOLD fluctuations in terms of decreasing E/I ratios, especially in higher-order association cortices [19]. Entirely different methods (machine learning-based inference) reveal that the effect of development on functional connectivity, especially in higher-order association areas, mimics the effect of a positive allosteric modulator increasing the efficacy of GABA-A receptors [18]. Thus, our results converge across a diversity of large-scale methods on the hypothesis that brain development through adolescence results in increased inhibitory versus excitatory drive in the mechanisms which generate and propagate θ and α oscillations.

Importantly, our results go beyond prior studies by suggesting not only a change in the E/I ratio, but separable effects on inhibition and excitation. Namely, considering effects on both E/I_HLP_ and E+I_HLS_, we infer that development results in an increase in inhibitory signaling (at least relative to excitatory signaling) in mechanisms driving θ-through-α waves (Figures 2,3,5). A relative increase in inhibitory signaling is consistent with enhanced GABAergic transmission in prefrontal cortex driven by a developmental increase in long-range glutamatergic inputs to parvalbumin (PV)-positive, fast-spiking GABAergic interneurons [9,10] and the increased expression of GABA α1 receptors [12] . This developmental increase in glutamatergic inputs may increase inhibitory signaling by promoting interneuron activity directly, and by upregulating PV genes from adolescence into adulthood, resulting in a more sustained boost in GABAergic signaling capacity.

While E/I decreases in low frequency bands, it increases in high-β and γ. Because γ rhythms are determined principally by the time constants of GABA-A receptor-mediated inhibition [37], increases in E/I in higher frequency bands may also, paradoxically, reflect increased GABA-A signaling. Greater signaling capacity, especially at fast GABA α1 receptors, can potentiate γ oscillations in response to excitatory perturbations by amplifying the alternating dynamics in reciprocally-connected inhibitory interneurons, or reciprocally-connected excitatory and inhibitory interneurons [37]. One consequence of these developmental changes would be an increase in spontaneous γ power. Indeed, we find that both E/I indices which scale monotonically with mean amplitude (fE/I and E/I_HLP_) increase with development in the γ band.

The combination of changes in critical dynamics and E/I imply that the systems driving specific frequencies shift differentially with respect to criticality. The strengthening of long-range temporal correlations and amplitude bistability in θ-through-β bands imply that these systems all shift towards criticality with development. In θ and α especially, the apparent decrease in E/I implies that these systems operate in a slightly super-critical regime in childhood and move towards criticality or even to a slightly sub-critical regime in adulthood. Conversely, the combination of weaker critical dynamics and an increase in E/I in the γ band for adults tells a different story. In terms of the fE/I index, γ oscillatory systems are sub-critical in children (fE/I << 1.0) and approach criticality (fE/I = 1.0) in adulthood (Figure 2C).

If γ systems shift towards criticality, why, then, do emergent dynamics become weaker? It is important to note that emergent dynamics of a critical system – like long-range temporal correlations and bistability – reflect spontaneous, endogenous activity patterns that can be disrupted by external perturbation [38]. Inhibitory suppression by θ or α waves, for example, might halt on-going γ oscillations, thus blunting long-range temporal correlations. This interpretation is consistent with the hypothesis that low-frequency oscillations exert top-down control over high-frequency oscillations [39–42] and is supported by evidence of an increasing propensity for cross-frequency coupling (e.g. δ–γ coupling: [43] and α–β coupling: [44]) across adolescent development. Moreover, our own analyses of branching ratios as a function of band-specific fE/I also suggest that cross-frequency interactions impact on spontaneous, endogenous activity. Namely, we find that greater excitability in high-frequency systems is predictably associated with faster growing avalanches, but the opposite is true of greater excitability in the systems that generate θ-band activity. One interpretation is that when θ-generating systems are more excitable, they suppress spontaneous, high amplitude activity endogenous to high-frequency systems. Such interactions could thus blunt long-range temporal correlations and bistability in γ-band activity that would otherwise emerge at criticality.

In addition to developmental effects on E/I and critical dynamics, we also observe state-dependent changes in the contrast of eyes-closed and eyes-open conditions (Figure 4). Specifically, opening eyes decreases E/I indices including both decreased fE/I and branching statistics. Consistent with reduced broadband power when eyes open [42,45], we find decreases in E/I ratios for all frequencies except the θ band where opening eyes increases E/I. We speculate that decreases in E/I reflect an increase in feedforward and feedback inhibition, to stabilize processing of visual inputs [46,47] when eyes open. Such effects may also reflect functional re-configurations of E/I to accommodate attentional demands.

This state manipulation also offers an opportunity to test the hypothesis that emergent dynamics are strongest when excitation and inhibition are balanced at criticality [48]. Namely, if opening eyes shifts brains in a subcritical direction, then individuals with supercritical dynamics (fE/I > 1.0) should show stronger long-range temporal correlations when they open their eyes, while those with subcritical dynamics (fE/I < 1.0) should show weaker long-range temporal correlations when they open their eyes (Figure 4D). Our findings (Figure 4F) support this prediction while also, thereby, validating fE/I as an individual difference index of E/I balance, with respect to criticality.

The facts that slow oscillatory systems (frequencies below γ) tend to be subcritical, and that opening eyes decreases long-range temporal correlations for all slower frequencies (except θ; Figure 4H) implies that slow oscillatory systems become increasingly subcritical with the onset of visual input. Indeed, it is hypothesized that brains tend to operate in a slightly subcritical, inhibition-dominant regime to maximize responsiveness while also preventing runaway, chaotic dynamics in response to excitatory inputs [49,50]. Interestingly, the brain continues to shift in a subcritical direction as cognitive demands increase beyond perception to working memory, judgement, and decision-making tasks [47,51–53], suggesting that an adaptive shift towards subcriticality scales with cognitive load. Conversely, the strengthening of long-range temporal correlations in the γ band when eyes open, despite the concomitant decrease in fE/I, implies that some mechanisms other than shifting E/I balance must unblock emergent dynamics. Perhaps decreased power in slower frequency bands also decreases top-down, cross frequency coupling, thus freeing γ to more robustly express long-range temporal correlations. Given that γ waves are implicated in conveying bottom-up information [39,41,54], unblocking emergent dynamics when eyes are opened would be adaptive for visual perception, enhancing, e.g., susceptibility and dynamic range [2–8] for the perception of a diversity of visual stimuli. Finally, we find that state-dependent changes in critical dynamics are stronger in adults than in children. The change in α- and β-band long-range temporal correlations when opening eyes, for example, is robust for adults, but non-existent in the youngest participants (Figure 4G). Such patterns suggest the intriguing hypothesis that one outcome of development is an increased capacity for adults to shift brain criticality to adapt to state-dependent processing demands.

A key limitation of our work is the use of EEG which restricts spatially-resolved inferences about developmental effects. Nevertheless, the temporal resolution of EEG supports distinctions among mechanisms generating slow versus fast oscillations, and inhomogeneity in topographical effect maps further implies that developmental effects on brain criticality vary by brain region. Future work using MEG, MRI, and invasive neural recordings will be better positioned to assess the role of developmental effects across distinct brain regions.

## Conclusion

Collectively, our results support the hypotheses that brains operate closer to criticality with development, and that this shift reflects mechanism-specific changes in E/I. We show that these developmental effects can also be evaluated in terms of emergent dynamics, as a function of neuronal oscillations, across frequency bands. Considering substantial evidence that criticality sets the stage for cognition by giving rise to emergent dynamics that support information processing, developmental shifts with respect to criticality have profound implications for cognitive function. Thus, our results motivate future studies examining how developmental shifts in emergent dynamics account for the enhancement of cognition across adolescent development and how brains adapt, across time, across state, and by mechanism, to accommodate to task demands.

## Methods

### Participants

N = 310 resting EEG recordings were collected from 164 healthy human participants (87/77 F/M) ranging in age from 10 to 33 years old at the University Pittsburgh Medical Center as part of a larger, longitudinal study characterizing developmental changes in signal processing supporting cognitive development. In an accelerated cohort design, participants each attended up to 3 visits, approximately 18 months apart. In addition to EEG and saccade task behavior, participants also completed a 7T MRI scan and a Spatial Span task (not analyzed here). Participants were recruited from the Greater Pittsburgh area and were excluded if they had a history of lost consciousness due to head injury, substance abuse, or major psychiatric or neurological condition in themselves or a first-degree relative, any learning disability, and non-correctable vision problems. All participants gave informed, written consent to participate or parents gave consent for those under 18 years old who provided assent. All experimental procedures followed an IRB-approved protocol, in compliance with the Declaration of Helsinki Code of Ethics of the World Medical Association.

### EEG Recordings and Preprocessing

Participants were seated upright in a sound-attenuated, electromagnetically-shielded room for EEG recordings using a 64-channel cap with passive electrodes and a Biosemi ActiveTwo amplifier. 1024 Hz recordings were collected during interleaved eyes-open and eyes-closed rest (four, one-minute intervals of each) and down sampled to 512 Hz. Additionally, data were high-pass filtered at 1 Hz using a Hamming windowed sinc FIR filter (using the *pop_eegfiltnew* function in EEGLAB), bad channels were removed (*clean_artifacts*) and interpolated (*pop_interp*), data were re-referenced to the average (*pop_reref*), and finally ICA was used (*pop_runica*) to identify components which, according to the ICLabel algorithm, were likely to be muscle, eye, heart, line noise, or channel noise artifacts.

### Measures of Brain Criticality and E/I Ratios

Emergent properties of a system at criticality include long-range temporal correlations, scale-free behavior, and multi-stability. We thus infer proximity to criticality from the strength of these properties.

Long-range temporal correlations refers to the propensity for the effects of a perturbation to persist longer than expected, e.g. by typical exponential decay. To measure *long-range temporal correlations* in EEG data, we use detrended fluctuation analysis (DFA) [26,27] to evaluate the persistence of amplitude fluctuations over time. Specifically, we extract the amplitude envelope via the Hilbert transform and take its cumulative sum over time. Next, the cumulative sum is broken up into windows of increasing duration. For each window size, the signal was broken into windows with 50% overlap, the linear trend is removed from each window, and the standard deviation of the detrended signal (referred to as the *fluctuation*) is computed. In systems with long-range correlations, this fluctuation function should grow rapidly as window durations increase. The rate of growth is quantified by the slope of a linear fit to a log-log plot of the mean fluctuation by window size. Higher slope values (closer to 1.0) indicate that the brain is operating closer to criticality [23].

Bistable dynamics arise when a system operates at the boundary of dynamical regimes because the system can express the dynamics of these distinct regimes over time without any further changes in control parameters. One clear example of bistability in EEG data is tendency of α waves to spontaneously transition between high and low amplitude oscillations over time [25]. The result of such transitions is that the amplitude distribution is better characterized by two modes rather than one. Here, following [28] we characterize bistability by the logarithm of the difference in the Bayesian Information Criteria scores for the fit of a bi-exponential versus a single exponential function to frequency-specific amplitude distributions. Higher values indicate that a bi-exponential model is an increasingly better description of the data and thus indexes criticality. First, we normalized the power time series (*P*) by its median value (converting *P* to *nP*), then we computed the probability density function by histogram (with 200 equally-spaced bins), and fit either a single, or bi-exponential function to the pdf using maximum likelihood estimation. The form of the single exponential model is

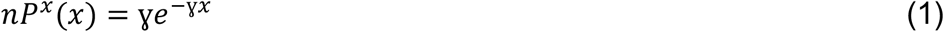

and the bi-exponential is

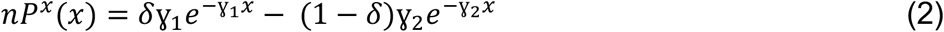

where γ_1_ and γ_2_ are exponents and δ weights the prevalence of each exponential in the overall pdf. Finally, we computed the BIC scores for each fit, correcting for differences in signal length and computed bistability (BiS) as the logarithm of the difference in BIC scores between the exponential and bi-exponential model.

E/I can also be characterized with respect to emergent dynamics. For example, the functional excitation-inhibition index (or fE/I), was developed to capture the empirical relation between long-range temporal correlations and signal amplitude in the vicinity of criticality [30]. Namely, in the slightly subcritical domain when inhibition is dominant, long-range temporal correlations and amplitude correlate positively, and in the slightly supercritical domain when excitation is dominant, this correlation is negative. fE/I is computed as

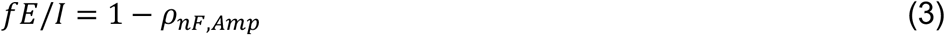

where *ρ_nF,Amp_* is the Pearson correlation between the normalized fluctuation function (*nF*) and amplitude and thus takes on a value of 1.0 at criticality. Here, following [30] we first bandpass filter our signal, then extract the amplitude envelope via the Hilbert transformation. Next, we compute the cumulative sum of the amplitude envelope and segment it into 80% overlapping, 5 s windows. Windows are then normalized by the mean amplitude in each window. Next, the linear trend in each window is removed and the fluctuation function is computed as the root-mean square of the residuals. The mean, normalized fluctuation function and mean amplitude are then correlated across windows to calculate fE/I. Note that because fE/I is only meaningful in proximity to the critical point, we only calculate fE/I values when the corresponding DFA value for that EEG time course is greater than 0.6.

Spontaneous bursts of neuronal activity are referred to as avalanches [55], and the growth rate of these avalanches can be used to infer the balance of excitation to inhibition. In EEG data, the growth rate of avalanches can be indexed in terms of the growth of bursts of high amplitude signal, clustered together in time [33,34,56]. Following [31] we estimate the branching statistic (instantaneous growth rate) by the following steps. First, we identify high amplitude bursts (or “events”) by thresholding EEG signal for each channel based on the point at which the amplitude distribution diverges from a best fit Gaussian model (2.5 SD in our data). Second, we discretize the timeseries by breaking into bins of width Δt. Here we chose Δt = .8 ∗ *IEI*, approximating each individual participants’ mean inter-event interval (IEI). We chose to use 80% of each participant’s mean IEI because that bin width maximized the correlation between fE/I and branching statistics across participants (cf. Figure 4A). Of note, results remain the same if we use 100% of each participants’ mean IEI or a fixed bin width across all participants (e.g. of 4 times the sampling rate, cf. [31]). Third, we identify avalanches as all events within contiguous time bins containing at least one event for any channel, separated by at least one time bin with no events. Fourth, we compute branching statistics as the ratio of the number of events in the second time bin of a given avalanche to the number of events in the first time-bin of a given avalanche. Finally, we compute a grand mean score by averaging branching statistics across all avalanches.

Another means of assessing E/I ratios in EEG data is to examine the ratio with which frequency-specific oscillations are high-amplitude versus low-amplitude, in a bistable system. Following [28] we use the fitted bi-exponential model parameter δ from Eqn. 4 which quantifies the weight of the pdf assigned to the high versus low power distribution in the bi-exponential model. This parameter, referred to as the proportion of high-to low-power oscillations index (E/I_HLP_) has been shown to index the strength of recurrent excitatory versus inhibitory drive (the E/I ratio) in prior work simulating oscillatory dynamics with a neural mass model [28]. While E/I_HLP_ can, in principle, take values from δ = 0 to 1, E/I_HLP_ is only interpretable in systems with bistable power dynamics. As such, we only compute E/I_HLP_ if BiS > 2.5 and only interpret values between 0.02 and 0.98 (c.f. Supplemental Figure S1).

Finally, we can also infer the combined strength of excitation and inhibition based on the separation between high- and low-power distributions. Indeed, for a given E/I ratio, the distance in amplitude separating the high- and low-power distribution grows as the sum of excitatory and inhibitory drive increases. Thus, the high-to-low-power separation index (E+I_HLS_) was developed to quantify summed neurotransmitter capacity. Following [28] we calculate E+I_HLS_ as the difference in the logarithm of the difference between the peak of the high-power exponential distribution and the peak of the low-power exponential distribution. As with E/I_HLP_, we only compute E+I_HLS_ if BiS > 2.5 and only interpret values of E+I_HLS_ when E/I_HLP_ is between 0.02 and 0.98.

### CROS Model Simulations

To simulate developmental effects on EEG dynamics, we adopted the CRitical OScillations model (from [23]), comprised of a 50 × 50 network of coupled, recurrent excitatory and inhibitory neurons that undergoes a critical phase transition as a function of the local connection densities of excitatory and inhibitory units. The CROS model generates oscillatory dynamics which evince criticality, including long-range temporal correlations and amplitude bistability when excitatory and inhibitory connectivity is balanced.

For our simulations, we used the same parameter set employed by Avramiea et al. (2025)[28], including 75% excitatory and 25% inhibitory neurons across our network, with excitatory and inhibitory connectivity ranging from 25% to 100% of neurons (in 2% steps across models) in a local grid of 7 × 7 neurons. 5 simulations for each combination of connectivity values were allowed to run for 1000 s. Net activity was calculated by summing up all spiking neurons at each time step, and adding in Gaussian white noise (μ = 0; σ = 3) to induce time-varying phase, which otherwise does not occur during silent periods in the network. All analyses of E/I_HLP_, and E+I_HLS_ were performed on the amplitude of this summed signal, for each simulation, and statistics were averaged across the 5 simulations of each combination of excitatory and inhibitory connectivity values.

