## Supplemental Figure 1 for "Brain criticality emerges with developmental shifts in frequency-specific excitation-inhibition balance"

**Supplemental Materials**

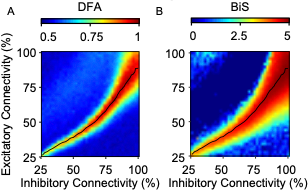

**Supplemental Figure 1** A) Long-range temporal correlations and B) bistability computed from amplitude fluctuations in CRitical OScillations model (CROS; Poil et al., 2012) simulations. Figures match the same set of simulations, across the same range of local excitatory and inhibitory connectivity presented in the main text (Figure 5). Figures reveal that emergent dynamics peak along the same line of excitation-inhibition balance as predicted for a critical phase transition.
